# Pantothenate auxotrophy in a naturally occurring biocontrol yeast

**DOI:** 10.1101/2022.12.14.519733

**Authors:** Maria Paula Rueda-Mejia, Raúl A. Ortiz-Merino, Stefanie Lutz, Christian H. Ahrens, Markus Künzler, Florian M. Freimoser

**Author notes:** Florian M. Freimoser, **Email**. **Author Contributions:** MPRM: investigation, data analysis, writing; RAOM: genome annotation; SL: genome assembly; CHA: genome assembly, review & editing; MK: review & editing, supervision; FMF: writing, supervision, project administration, funding acquisition. **Competing Interest Statement:** The authors have no competing interests.

## Abstract

The genus *Hanseniaspora* is characterized by some of the smallest genomes among budding yeasts. These fungi are primarily found on plant surfaces and in fermented products and represent promising biocontrol agents against notorious fungal plant pathogens. In this work, we identify a *Hanseniaspora meyeri* isolate that shows strong antagonism against the plant pathogen *Fusarium oxysporum* as a pantothenate auxotroph. Furthermore, strong biocontrol activity *in vitro* required both pantothenate and biotin in the growth medium. We show that the *H. meyeri* isolate APC 12.1 can obtain the vitamin from plants and other fungi. The underlying reason for the auxotrophy is the lack of key pantothenate biosynthesis genes, but at least six genes encoding putative pantothenate transporters are present in the genome. By constructing and using a *Saccharomyces cerevisiae* reporter strain, we identified one *Hanseniaspora* transporter, out of the six candidate proteins, that conferred pantothenate uptake activity to *S. cerevisiae*. Pantothenate auxotrophy is rare and has only been described in a few bacteria and in *S. cerevisiae* strains that were isolated from sake. Such auxotrophic strains may seem an unexpected and unlikely choice as potential biocontrol agents, but they may be particularly competitive in their ecological niche and their specific growth requirements are an inherent biocontainment strategy preventing uncontrolled growth in the environment. Auxotrophic strains such as the *H. meyeri* isolate APC 12.1 may thus represent a new strategy for developing biocontrol agents that will be easier to register than prototrophic strains, which are normally used for such applications.

**Significance Statement:** As a precursor of the essential coenzyme CoA, pantothenate is present in all organisms. Plants, bacteria and fungi are known to synthesize this vitamin, while animals must obtain it through their diet. Pantothenate auxotrophy has not been described in naturally occurring, environmental fungi and is an unexpected property for an antagonistic yeast. Here, we report that yeasts from the genus *Hanseniaspora* lack key enzymes for pantothenate biosynthesis and identify a transporter responsible for the acquisition of pantothenate from the environment. *Hanseniaspora* isolates are strong antagonists of fungal plant pathogens. Their pantothenate auxotrophy is a natural biocontainment feature that could make such isolates interesting candidates for new biocontrol approaches and allow easier registration as plant protection agents compared to prototrophic strains.

## Introduction

The genus *Hanseniaspora*, the teleomorph of *Kloeckera*, comprises widely distributed apiculate yeasts that are among the most abundant fungi on fruits and reach high cell densities during the early stages of fermentation in wine must (1–3). Several *Hanseniaspora* strains show potential as biocontrol agents against plant pathogens of important fruit crops. Antagonistic activity against crop spoilage moulds such as *Botrytis, Corynespora, Rhizopus, Penicillium*, or *Phytophthora* has been described in grape, apple, citrus and strawberry fruits (4–8).

Growth assays with *Hanseniaspora* culture filtrates, supernatants and sterilized solutions did not result in mould inhibition (9). Therefore, the mechanism for antagonism was presumed to be competition for nutrients and space. However, the interest in a mechanistic understanding of biocontrol activities has grown recently and other possible mechanisms may be involved in the *Hanseniaspora* activity against plant pathogens. Inhibition of *Botrytis cinerea* spore germination and mycelium growth by *Hanseniaspora uvarum* has been described (5). Volatile and soluble compounds were also found to be involved in the biocontrol activity of *Hanseniaspora*. Specifically, volatile organic compounds (VOCs) such as 1,3,5,7-cyclooctatetraene are produced by *H. uvarum* in the presence of *B. cinerea* on strawberry surfaces, and compounds extracted from *H. osmophila* had significant inhibitory effects on the same pathogen (10, 11).

The genus *Hanseniaspora* displays some of the smallest genomes and sets of annotated genes among the budding yeasts in the order Saccharomycetales (on average 9.71 ± 1.32 Mbp and 4708 ± 634 genes)(12). The *Hanseniaspora* clade has lost a large number of genes encoding proteins with functions in primary metabolism, cell-cycle regulation and the maintenance of genome integrity. Due to such gene losses, many *Hanseniaspora* strains are unable to metabolise certain sugars (e.g., galactose, maltose, raffinose, or melezitose), are auxotrophic for thiamine or specific amino acids, and exhibit accelerated evolution due to their high mutation rate and genome instability (12). Considering their small genomes and limited metabolic capabilities, the success in various environmental habitats and strong competitive activity against pathogenic fungi *in vitro* is remarkable and raises the question about the underlying mechanisms.

Previously, we reported a *Hanseniaspora* meyeri isolate from apple flowers that showed strong antagonistic activity against a wide variety of important plant pathogens in agar-based competition assays (13). Here, we show that this isolate is a pantothenate auxotroph and can obtain this essential vitamin from plant roots and filamentous fungi. While the *Hanseniaspora* strain APC 12.1 (as well as other strains of this genus for which whole genome sequences are available) lacks genes for pantothenate biosynthesis, its genome contains at least six candidate pantothenate transporter genes (based on KEGG annotations). One of these transporters, named Pant4, was shown to complement the phenotype of a *S. cerevisiae* strain lacking its endogenous pantothenate symporter Fen2, encoded by a single copy gene.

In summary, these studies describe an antagonistic, environmental yeast with a reduced genome that lacks the ability to synthesize pantothenate, but that can obtain this vitamin from plants and fungi in its ecological niche via at least one pantothenate transporter. Such strains are interesting for novel biocontrol applications, because the pantothenate auxotrophy prevents uncontrolled spread in the field, but can easily be overcome by adding the vitamin to the formulation.

## Results

### On plant growth medium, *H. meyeri* (APC 12.1) must obtain an essential nutrient from plant roots or other fungi

In the past, we have identified several yeasts that strongly inhibit filamentous fungi in binary competition assays *in vitro* (13). Here, the goal was to assess yeast biocontrol activity in the presence of a host plant. The assay comprised the yeast *H. meyeri* (isolate APC 12.1), the plant pathogen *Fusarium oxysporum* f. sp. *lycopersici* (FOL), and tomato seedlings and was performed on plant culture medium lacking a carbon source (to prevent overgrowth by the yeast and FOL).

In these assays, *H. meyeri* (APC 12.1) only grew around the FOL colony and, to some degree, in the vicinity of the tomato root (Figure 1A). In contrast, this yeast grew on the entire plate surface during competition assays on complete YNB and PDA media (Supplementary Figure 1). This suggests that the yeast must obtain an essential nutrient from the fungal colony or the plant root when plated on Murashige and Skoog (MS) medium that is used for the *in vitro* cultivation of plants. Independent *Hanseniaspora* growth tests were performed with the same medium and with radish seedlings, or with the filamentous fungi *Penicillium polonicum, Mucor moelleri*, or *Botrytis caroliniana*. In all assays, the same yeast growth pattern was observed: *H. meyeri* colonies were growing in the vicinity of a competing fungus or a host plant and no colonies were observed further away from either organism (Supplementary Figure 2). These observations suggested that, on MS medium, *H. meyeri* obtains an essential nutrient from competing fungi or from plants.

**Figure 1.**
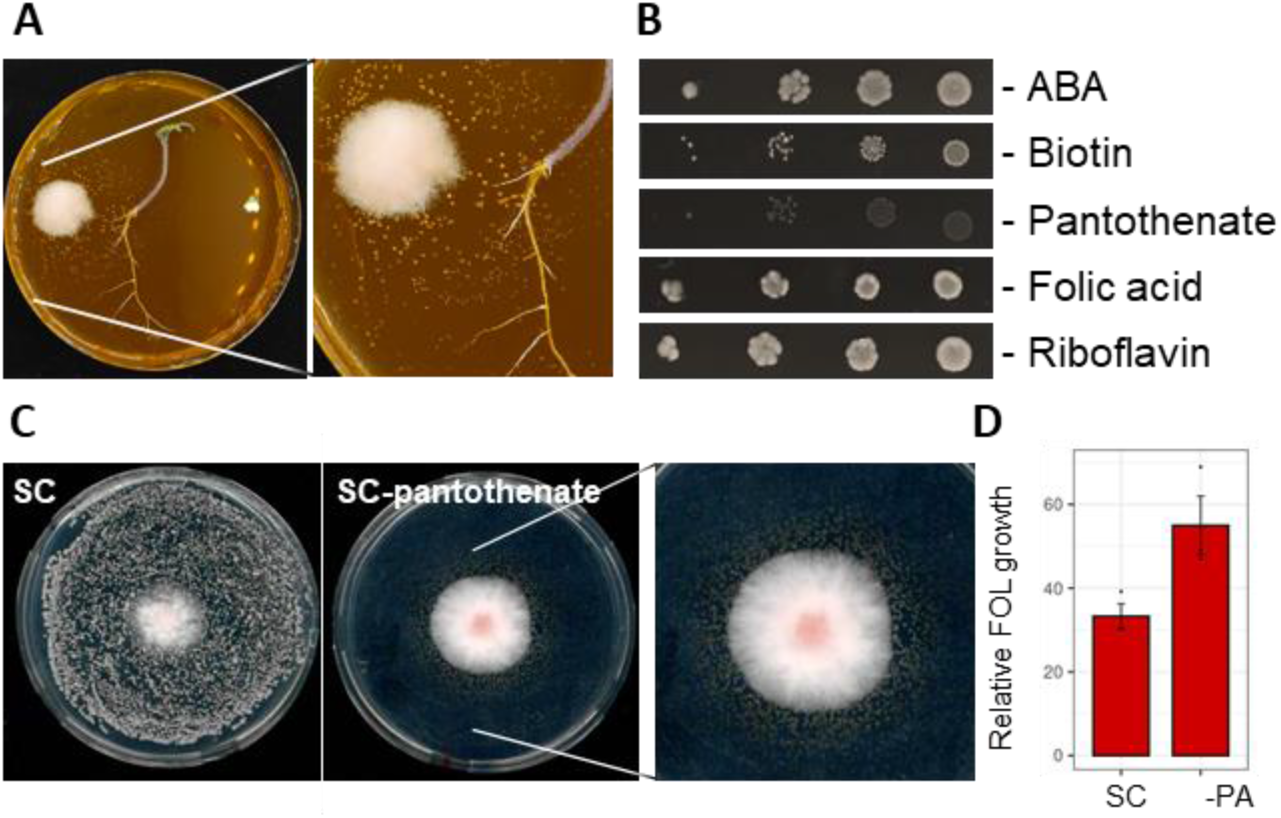
*Hanseniaspora meyeri* (APC 12.1) is auxotrophic for pantothenate. **A**. Tomato-*F. oxysporum*-yeast interaction on MS medium. **B**. *H. meyeri* (APC 12.1) growth on SC medium lacking amino benzoic acid (ABA), biotin, calcium pantothenate, folic acid, or riboflavin. *H. meyeri* was unable to grow in the absence of calcium pantothenate. The lack of either biotin or folic acid resulted in reduced growth and colony size. Medium without riboflavin or ABA permitted normal growth. **C**. *Fusarium-Hanseniaspora* competition in SC and SC without pantothenate. **D**. On SC medium lacking calcium pantothenate *H. meyeri* only grew around the FOL colony and inhibited growth. The relative growth, as compared to the FOL colony area in the absence of a yeast, was reduced to 55% in SC lacking pantothenate (-PA) and to 33% in complete SC medium.

### *H. meyeri* (APC 12.1) is auxotrophic for pantothenate

In order to identify the essential nutrient(s) that *H. meyeri* obtains from fungi or plants, growth experiments with different defined media were performed. While the yeast did not grow on MS medium, it grew well on a synthetic complete medium (SC) containing yeast nitrogen base (YNB) with a complete supplement mix. MS medium lacks different vitamins that are present in standard microbiological culture media. To identify which vitamin the *H. meyeri* strain APC 12.1 obtains from fungi or plants, we performed growth assays with amino benzoic acid (ABA), biotin, calcium pantothenate, folic acid, and riboflavin.

*H. meyeri* grew well in the absence of either ABA or riboflavin, showed reduced growth in media lacking biotin or folic acid, but was unable to grow in the absence of calcium pantothenate (Figure 1B). Similarly, *H. meyeri* growth on MS medium could be recovered by adding calcium pantothenate, though colonies were relatively small. On SC medium lacking pantothenate, *H. meyeri* only grew in the vicinity of the FOL colony, as was observed on MS medium (Figure 1C). The *Hanseniaspora* cells growing around FOL inhibited *Fusarium* growth to 55% (as compared to growth in the absence of any yeast), while on complete SC medium the *Fusarium* colony was reduced to 33% of the control (Figure 1D). These findings suggested that *H. meyeri* (APC 12.1) is unable to synthesize pantothenate and must obtain this vitamin from the environment.

### Biotin and calcium pantothenate are required for biocontrol activity of *Hanseniaspora* against *Fusarium*

To test the effect of different vitamins on the biocontrol activity of *H. meyeri* against *Fusarium*, competition assays on different MS and SC media were performed. FOL growth was not inhibited by the presence of *H. meyeri* on SC media that specifically lacked either biotin or folic acid (Figure 2A). However, *H. meyeri* inhibited FOL growth in the absence of ABA, calcium pantothenate, or riboflavin (as compared to growth in the absence of the yeast). In the presence of *H. meyeri*, FOL growth was reduced by more than 50% on complete SC medium and in the absence of ABA, while this inhibition was lower when either calcium pantothenate or riboflavin were omitted (Figure 2A). On MS medium, the addition of calcium pantothenate alone did not restore antagonistic activity of *H. meyeri* (Figure 2B). We therefore supplemented MS medium with pantothenate and each of the other four vitamins that were tested before and performed competition assays. Only the addition of pantothenate and biotin resulted in strong FOL inhibition by *H. meyeri* (over 40%) (Figure 2B). These two vitamins were sufficient to recover inhibitory activity to the same degree as the addition of all vitamins.

**Figure 2.**
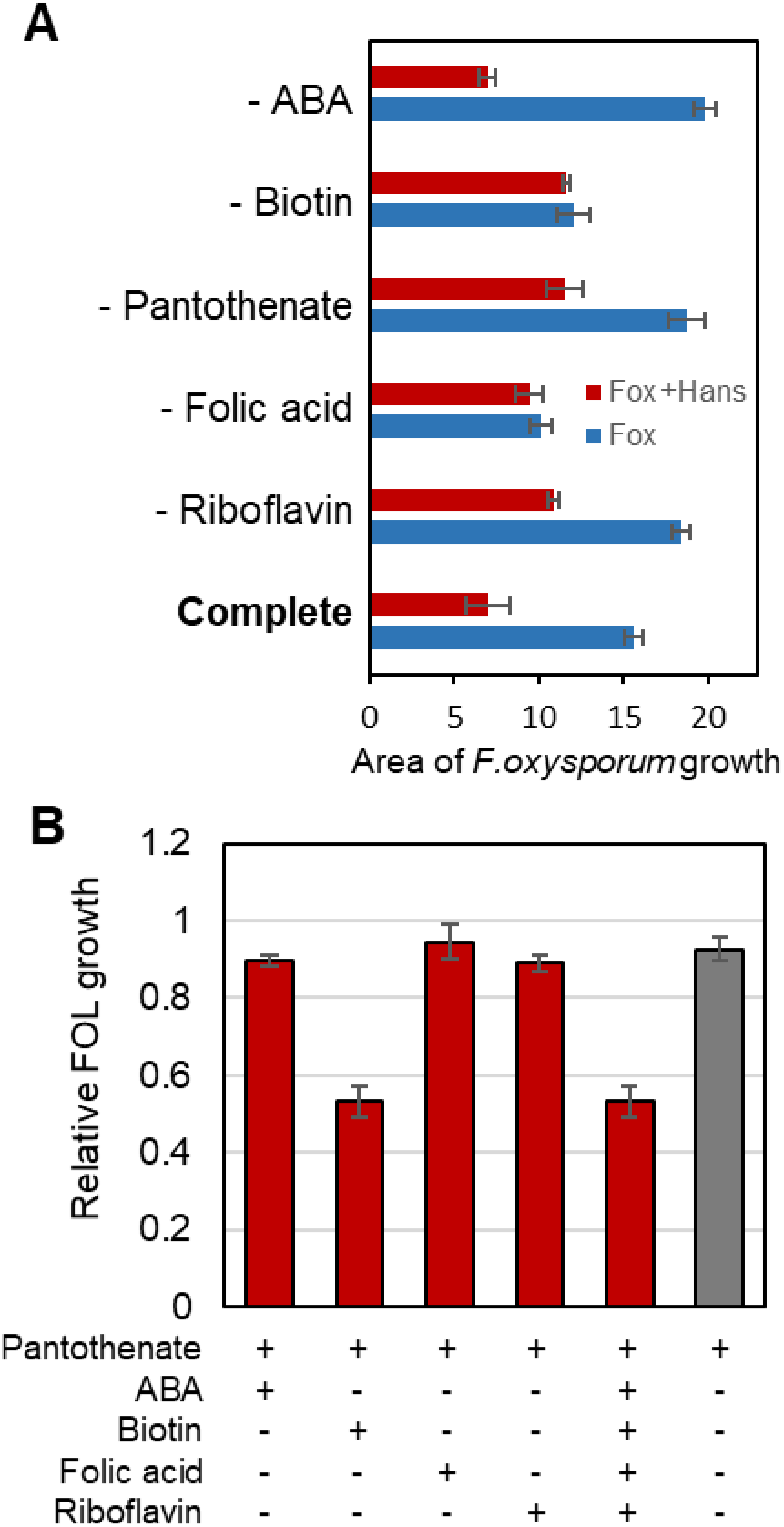
Biotin and calcium pantothenate are required for biocontrol activity of *Hanseniaspora* against *Fusarium*. **A**. Absence of vitamins in SC medium affects antagonistic activity. In complete SC medium or when ABA is lacking, *Hanseniaspora* strongly inhibits the growth of *Fusarium* (red) as compared to the control without the yeast (blue). The activity is reduced without calcium pantothenate or riboflavin and abolished in the absence of either biotin or folic acid. **B**. Addition of biotin and calcium pantothenate to MS medium results in the same inhibitory activity as in the medium with the five vitamins. Supplementation with pantothenate and either one of the other vitamins did not improve inhibitory activity.

These results demonstrate that biotin and calcium pantothenate are required for *H. meyeri* biocontrol activity against FOL. Even though *H. meyeri* is auxotrophic for pantothenate, it inhibited FOL on SC medium lacking only calcium pantothenate. This yeast must thus possess effective transporters that can take up vitamin secreted from FOL mycelium.

### The *H. meyeri* (APC 12.1) genome lacks crucial genes associated with pantothenate biosynthesis

In order to identify the genetic basis for pantothenate auxotrophy and ability to grow with externally supplied pantothenate, the genome of *H. meyeri* (APC 12.1) was sequenced using a combination of long (Pacific Biosciences and Oxford Nanopore Technologies) and short (Illumina) reads, *de novo* assembled, and analyzed. The final genome assembly consisted of seven complete chromosomes and a 17 kb mitogenome (Supplementary Table S1). The latter showed inverted repeats, which suggest a linear mitochondrial genome as it has been observed in other yeasts (14– 16). In our genome assembly, on chromosome 2 (approximately at position 1060 kb), two 96% identical tandem rDNA units were present. Based on the YeastIP database, the first rDNA unit was 96% (ITS) and 95% (D1/D2 region) identical to *Hanseniaspora clermontiae* and *H. meyeri*, respectively, while the second unit showed 99% identity to the corresponding sequences from both species. The large *MDN1* gene (ORF: HANS 0B04920, at 950 kb on chromosome 2) was 99% identical to the corresponding gene from *H. meyeri*, but only 93% identical to *H. clermontiae*. All other genes that we tested were also most similar to the corresponding *H*. meyeri homolog. The isolate APC 12.1 is thus named and referred to as *H. meyeri*.

The pantothenate biosynthetic pathway starts with L-aspartate and L-valine as substrates and results in the formation pantoate and β-alanine, which are the substrates for the finial reaction catalysed by pantothenate synthase (Figure 3A). The *H. meyeri* APC 12.1 genome contained homologs of most genes involved in this pathway, but we could not detect a 3-methyl-2-oxobutanoate hydroxymethyltransferase gene (EC 2.1.2.11, *ECM31* in *S. cerevisiae*) and only a weak hit (E > 10^−10^) for pantothenate synthase (EC 6.3.2.1, *PAN6* in *S. cerevisiae*). A genome analysis of different Saccharomycetales species showed the absence of *PAN6* (EC 6.3.2.1) and *ECM31* (EC 2.1.2.11) homologs in all *Hanseniaspora* genomes available at Mycocosm, while all other selected yeasts (model organisms, human pathogens and yeasts with biotechnological or biocontrol potential) harbored genes corresponding to all the enzyme types upstream of pantothenate biosynthesis in *S. cerevisiae* (Figure 3B). The absence of these two genes makes it impossible for *H. meyeri*, and other isolates of this genus, to synthesize pantothenate and is thus the molecular cause for the observed pantothenate auxotrophy phenotype.

**Figure 3.**
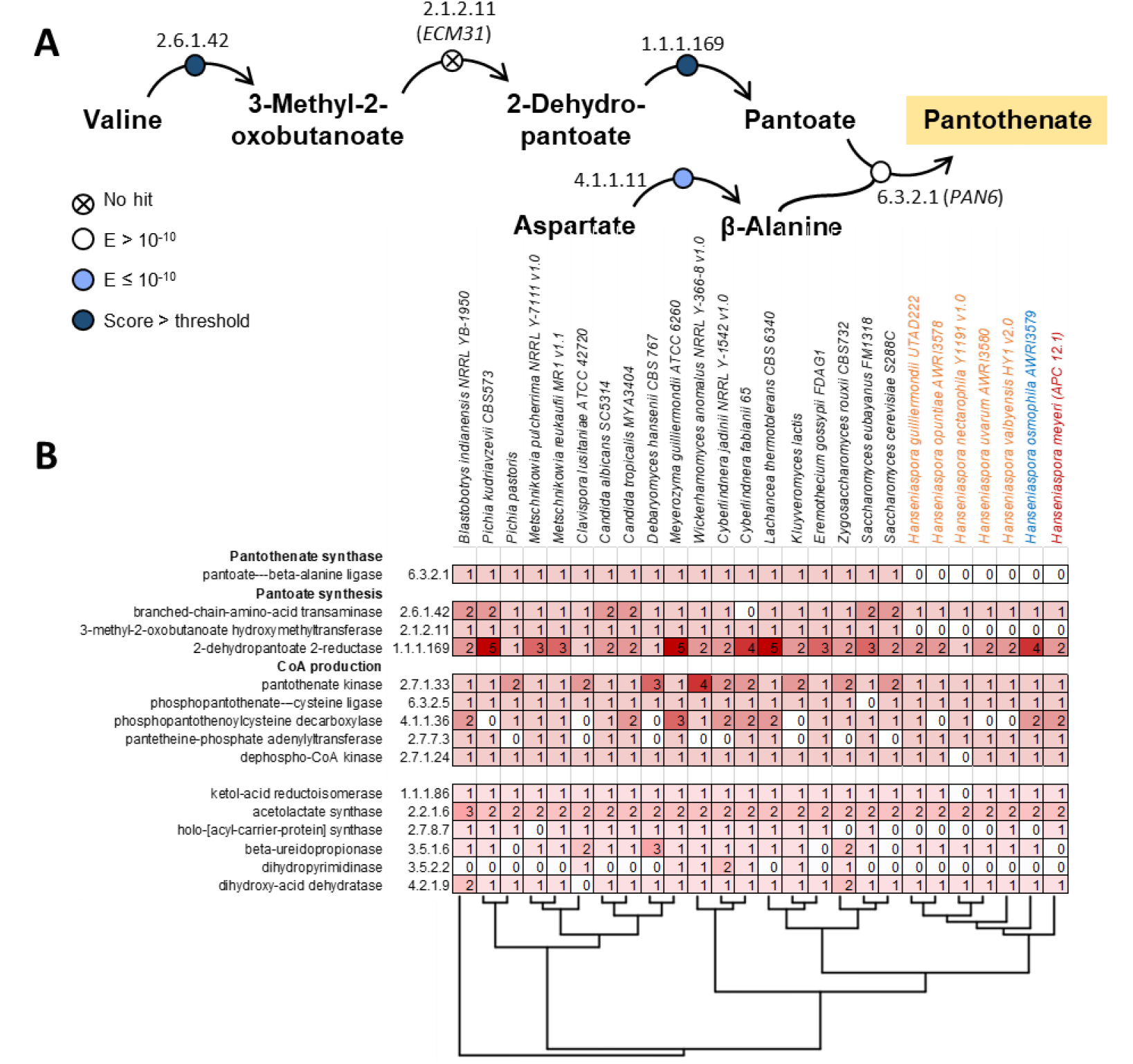
A. The *Hanseniaspora meyeri* (APC 12.1) genome lacks crucial genes associated with pantothenate biosynthesis. **A**. The pantothenate biosynthesis pathway from *S*.*cerevisiae* showing enzymes with a homolog in *H. meyeri*. The significance of the best BLAST hit is indicated in white (E> 10-10 or low significance), light blue (E ≤ 10-10) or dark blue (highly significant). **B**. Presence of genes in the pantothenate biosynthesis pathway in a selection of budding yeasts (Saccharomycotina). The number of genes per enzyme category is highlighted in shades of red (0 is white and 5 the darkest red). Selected representatives of the subphylum and six *Hanseniaspora* species were included along with the isolate APC 12.2. *Hanseniaspora* species from the fast evoving lineage are shown in orange, while slow evolving strains are depicted in blue (as described by Steenwyk et al. 2019). The tree corresponds to the phylogenetic relationships of the selected species. The mentioned absence of two key genes for pantothenate biosynthesis occurs in all *Hanseniaspora* representatives. Meanwhile, homologous genes for the steps upstream of pantothenate are found in all other yeasts except *Cyberlindnera fabianii*, which lacks the branched-chain-amino-acid transaminase (EC 2.6.1.42). Downstream from pantothenate, the CoA production pathway shows the absence of a variety of genes across the tree.

### The *H. meyeri* (APC 12.1) genome harbors at least six putative pantothenate transporter genes

Pantothenate is a crucial precursor of CoA and thus essential for various metabolic functions, including the biosynthesis of fatty acids and sterols in the citric acid cycle, and gene regulation through histone acetylation. A cell that is unable to synthesize pantothenate thus must possess transporters that allow uptake of this vitamin. In order to identify potential pantothenate transporters in *H. meyeri*, we performed genome analyses and searched KEGG annotations.

The KofamKOALA KEGG analysis of the annotated *H. meyeri* APC 12.1 genome predicted six genes that might encode pantothenate transporters (ORF numbers 0A02930, 0A05630, 0B05620, 0D01900, 0D03170, and 0E01180, respectively) (Figure 4). These genes were named *PANT1-6* and encoded proteins ranging in size from 419 to 466 amino acids. Structural predictions, based on the amino acid sequences, estimated all of these six proteins to be α-helical bundles. Except for Pant3 (14 transmembrane domains, N- and C-terminus facing the extracellular space), 12 transmembrane domains and cytosolic N- and C-termini were predicted for all proteins. Pant1, Pant2 and Pant5 were assigned to the major facilitator superfamily and the Glycerol-3-phosphate transporter family. The sole *S. cerevisiae* pantothenate transporter Fen2 also belongs to the major facilitator superfamily and, as most members of this superfamily, has 12 transmembrane helixes. However, it belongs to the allantoate permease family, a group of yeast transporters importing anionic vitamins (17, 18). Among the six putative *H. meyeri* pantothenate transporters, Pant4 had the highest similarity to Fen2 (50% amino acid identity).

**Figure 4.**
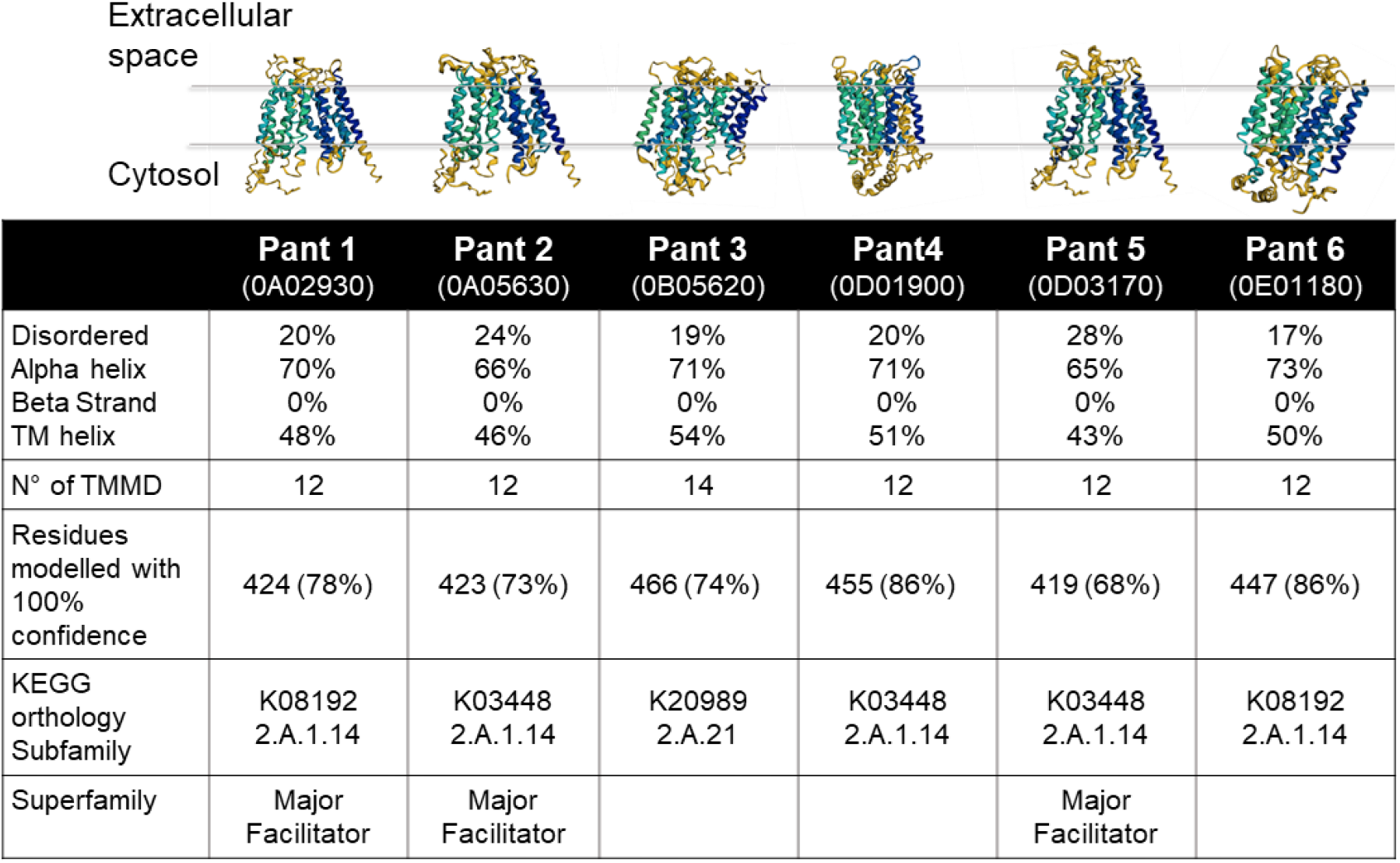
The *Hanseniaspora meyeri* (APC 12.1) genome harbors at least six putative pantothenate transporter genes. Amino acid sequences of the six transporters were analyzed with Phyre2. All six predicted proteins are formed by 65 to 73% alpha helixes, out of which the most part (43 to 54% of the total protein) form the transmembrane domains (TMMD). The number of transmembrane domains is predicted to be 12 for all the transporters except for Pant3, for which 14 such domains are proposed.

Based on genome analyses, we thus identified six *H. meyeri* (APC 12.1) genes as candidates for encoding a functional pantothenate transporter that enable this yeast to grow and compete in the environment.

### Complementation with *H. meyeri PANT4* restores pantothenate uptake in *S. cerevisiae*

*S. cerevisiae* encodes a single pantothenate transporter, Fen2, and is also able to synthesize the vitamin. However, if the vitamin is absent from glucose-containing media, growth is slightly impaired, suggesting that pantothenate uptake is limiting under these conditions. A *Δpan6* deletion mutant that lacks the enzyme (EC 6.3.2.1) required for the final step of pantothenate synthesis (Figure 3A) has a comparable phenotype as *H. meyeri* APC 12.1 (i.e., it does not grow without pantothenate but can obtain the vitamin from other fungi). In a *Δpan6* strain background, the Fen2 transporter is essential because pantothenate cannot be synthesized and must therefore be taken up. This strain can thus be taken advantage of to create a reporter strain for pantothenate transport. For this purpose, the *FEN2* promoter was exchanged with the galactose inducible promoter GALL and putative *Hanseniaspora* pantothenate transporters were heterologously expressed by integrating the corresponding genes at the *URA3* locus.

As expected, the reporter strain grew comparably to wild type yeast and showed no growth defect in YNB medium containing galactose (“control” in Figure 5A), while it was unable to grow with glucose as the sole carbon source. Constructs of the six predicted potential *H. meyeri* (APC 12.1) pantothenate transporter genes *PANT1-6* (untagged, 3xHis+6xFLAG-, or mRuby-tagged) were transferred and expressed under a constitutive promoter in this reporter strain. Western blot analysis confirmed the expression of three of the six transporters, Pant2, Pant4 and Pant5 (Supplementary Figure 3). Spot assays showed that out of the six tested transporter genes, only *PANT4* was able to recover growth of the reporter strain on glucose-containing medium, suggesting that Pant4 is a pantothenate transporter with a homologous function to *S. cerevisiae* Fen2. This result was the same for the cells harboring the tagged and untagged constructs. Confocal microscopy localized the mRuby-Pant4 fusion protein to the plasma membrane, as confirmed by colocalization with a green fluorescent plasma membrane dye (Figure 5B). Overall, these results show that Pant4 from the auxotrophic isolate *H. meyeri* APC 12.1 functions as a pantothenate transporter in *S. cerevisiae*.

**Figure 5.**
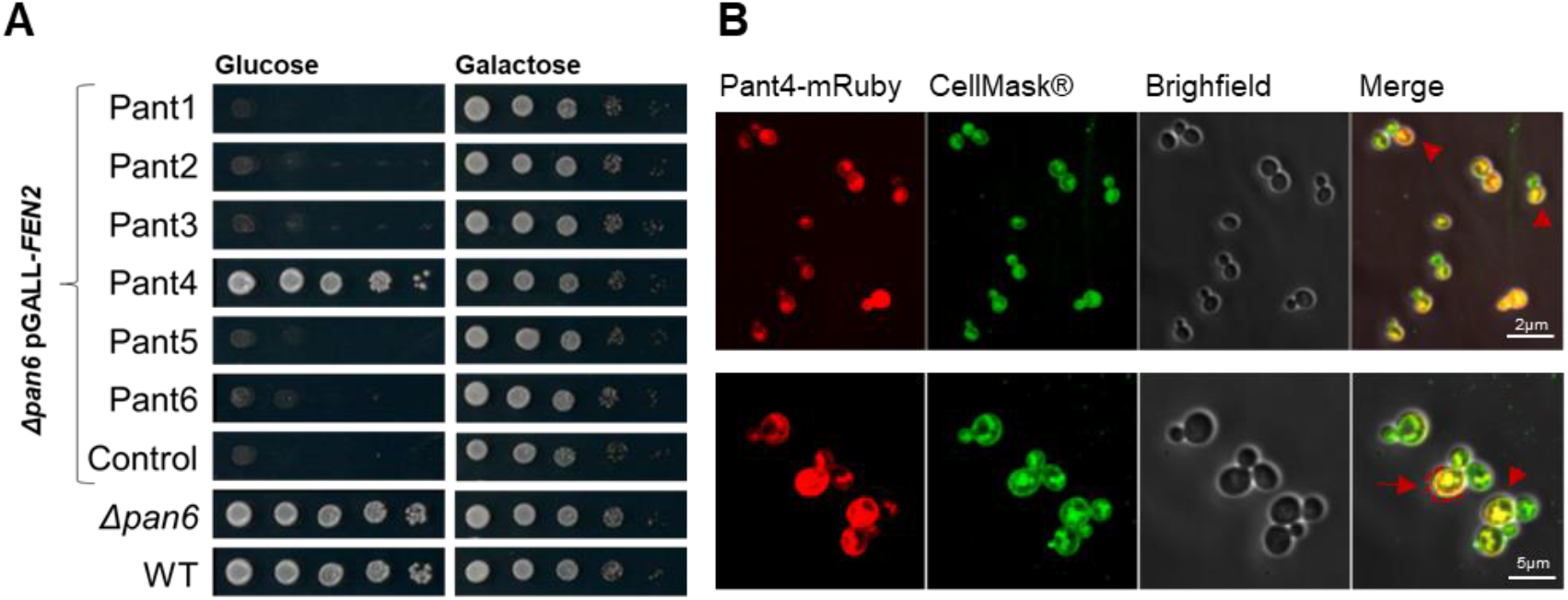
Complementation with *H. meyeri PANT4* restores pantothenate uptake in *S. cerevisiae*. Heterologous expression of *PANT4* restores growth in the *S. cerevisiae* reporter strain. **A**. *H. meyeri* predicted pantothenate transporters *PANT1-6* were constitutively expressed in the *Δpan6 pGALL-FEN2* reporter strain, which is unable to both produce pantothenate and obtain it from the medium when glucose is the carbon source. Out of the six transporters, only Pant4 expression is able to recover normal growth in glucose. **B**. Pant4 localizes to the plasma membrane of mother cells, as confirmed by its colocalization with the CellMask® green plasma membrane dye.

## Discussion

The study presented here describes the unexpected discovery of a naturally occurring, pantothenate auxotrophic *H. meyeri* (APC 12.1) strain that exhibits strong biocontrol activity against plant pathogens. Auxotrophic microorganisms might seem an unlikely choice for applications such as biocontrol because of their metabolic limitations. However, the use of such strains may constitute a natural biocontainment strategy and mitigate concerns about uncontrolled propagation in the environment. Natural auxotrophs should thus be easier to register as biocontrol agents, while their metabolic limitations can likely be overcome by strain-specific formulations. Interestingly, biocontainment strategies are actively studied and developed in order to prevent the spread of synthetic organisms (or “normal” genetically modified microorganisms) in the environment (19–21). Such tools are considered essential for possible applications of synthetic organisms and are thus of considerable economic interest (22, 23). Based on the observations provided here, specific formulations and application strategies that include calcium pantothenate, but also other vitamins such as biotin and folic acid, could be envisioned.

The *H. meyeri* (APC 12.1) genome lacks key genes for the biosynthesis of pantothenate, which is likely the root cause of the observed pantothenate auxotrophy. Such a pantothenate auxotrophy has also been described in the bacterial species *Zymomonas mobilis* and *Methylobacterium* spp. (24, 25). In these examples, auxotrophy was due to the lack of the corresponding aspartate decarboxylase PanD or one of the enzymes required to synthesize the precursor β-alanine, respectively. Interestingly, these bacteria are also found associated with plants and in fermentation processes, suggesting that these environments are able to sustain the growth of pantothenate auxotrophs. Similarly, pantothenate auxotrophy (and also auxotrophy for biotin, niacin, and thiamine) is frequent among bacteria living in the *Arabidopsis thaliana* phyllosphere (26). However, in yeasts pantothenate auxotrophy has only been described for *S. cerevisiae* strains isolated from sake and is due to a loss of function in the Ecm31 gene, which impedes the synthesis of pantoate (27). A recent study isolated 433 wild yeasts from a variety of raw and fermented cereals. All isolates, belonging to 9 species, grew in the absence of pantothenate. In contrast, biotin and riboflavin were necessary for the growth of 64% and 14.3% of all strains, respectively (28). Pantothenate auxotrophy is thus rare in nature and has not been described for environmental fungi. Although only few reports are available, secretion of pantothenate or pantothenate precursors seems a common phenomenon, as it has been described for *Arabidopsis thaliana*, root exudates from different plants, *Cryptococcus neoformans* (as a quorum sensing signal), or even infected lung epithelial cells (25, 29–31). Pantothenate therefore seems readily available in plant environments, which likely explains why this auxotrophy can arise in plant-associated yeasts and bacteria.

The genus *Hanseniaspora* is characterized by some of the smallest genome sizes and lowest gene numbers in budding yeasts. A genomic study with 25 genomes from 18 *Hanseniaspora* species described a faster- and a slower-evolving *Hanseniaspora* lineage and uncovered large-scale gene loss as a means of genome evolution in both (12). With these gene losses, a number of metabolic functions, as well as cell-cycle checkpoints, disappeared. Gene functions related to growth in and fermentation of maltose, degradation of sucrose, and galactose assimilation are often absent. Several *Hanseniaspora* species are thus not able to grow with maltose, raffinose, melezitose, galactose or sucrose as the sole carbon sources. Another vitamin, thiamine, is also not synthesized by some species, as most of the biosynthetic pathway has been lost. The species *H. meyeri* belongs to the faster-evolving *Hanseniaspora* lineage and is shown here to be metabolically restricted and dependent on cross-feeding of essential nutrients from plants and competing microorganisms. Although an auxotrophic phenotype may seem a competitive disadvantage, the loss of biosynthetic genes can confer a selective advantage to bacteria and stabilize microbial communities (32–36). In addition, the absence of components of the cell cycle and the reduced genome size are thought to accelerate cell division and growth, explaining the high cell densities rapidly reached in fermentation products. The combination of these two adaptive processes could thus explain the strong biocontrol activity we observed for the *H. meyeri* APC 12.1 strain. Rapid cell growth and colonization, combined with the conservation of energy due to the uptake, instead of the costly biosynthesis, of essential vitamins, may give *H. meyeri* the competitive advantage necessary to antagonize pathogenic fungi. This might thus represent a prime example for the “competition for nutrients” mode of action that is, counterintuitively, based on and made possible by vitamin auxotrophy.

In general, competition for nutrients and space has been suggested as the main mechanism of yeast biocontrol against fungal pathogens (37). Yeasts grow faster and colonize the available space, take up and deplete essential nutrients, and reduce pathogen growth (36). However, few studies describe this phenomenon in detail or at a molecular level (37). Here, we describe auxotrophy for the vitamin pantothenate, which may eventually allow the yeast to compete more efficiently for other nutrients and impede the growth of plant pathogenic fungi. Auxotrophies, such as the one described here, and the high competitiveness in general, could also be taken advantage of for biocontrol applications in agriculture. Including nutrients and vitamins in the formulation of biocontrol yeasts could improve the establishment and persistence of these organisms on the crop and thus result in a more reliable biocontrol activity. Detailed analyses of the metabolic capabilities and uptake efficiencies for different nutrients may thus greatly benefit biocontrol and lead to new and more successful applications of antagonistic yeasts in crop protection.

## Materials and Methods

### Strains and cultivation

All strains used in this study are listed in Supplementary Table S2. *H. meyeri* (APC 12.1) was isolated from apple flowers in Switzerland (13). For competition assays, *Fusarium oxysporum* f. sp. *lycopercisi*, kindly provided by Antonio Di Pietro, was used. Tomato seeds (cultivar Moneymaker) were surface sterilized and germinated on Murashige and Skoog (MS) agar. The *S. cerevisiae Δpan6* deletion strain was obtained from EUROSCARF.

Interaction assays with *H. meyeri, F. oxysporum* and tomato seedlings were performed on MS (Duchefa Biochemie, Haarlem, Netherlands). Overnight *H. meyeri* cultures were collected, the pellet was washed with water, and 100 μL of yeast solution (OD_600_ ≈ 0.1) was spread on the plate. *F. oxysporum* conidia were isolated after growth for 3-4 d in potato dextrose broth (PDB; Becton, Dickinson and Company, Le Pont de Claix, France) by filtering through two layers of Miracloth^®^ (Merck Millipore, Schaffhausen, Switzerland). Conidia were collected and washed twice with water, the concentration was estimated by hemocytometer counting, and solutions with 5 × 10^5^ conidia/mL were prepared. A 5 μL drop of conidia solution was placed on one side of the plate and one week-old plants were placed on the other side. Competition assays with *H. meyeri* and *F. oxysporum* in the presence or absence of different vitamins were performed with SC medium (Formedium™, Norfolk, United Kingdom) and as described previously (13).

### Genome sequencing and annotation

*H. meyeri* (APC 12.1) genomic DNA was extracted using a phenol/chloroform extraction protocol and as previously stated (38, 39). Cells from an overnight culture were collected and resuspended in 200 μL of Harju Buffer (2% Triton X-100, 1% SDS, 100 mM NaCl, 10 mM Tris-HCl, pH 8.0, 1 mM EDTA). Disruption was performed by two rounds of freezing with liquid nitrogen and boiling at 95°C. A high coverage genome was produced using a combination of Oxford nanopore, Illumina and PacBio technologies. Libraries were prepared using the Nanopore ligation sequencing kit and the Illumina Nextera XT DNA library prep kit. After filtering the PacBio and Oxford Nanopore subreads with Fitlong (v.0.2.0), and the Illumina reads with trimmomatic (v0.39), we performed a *de novo* assembly of PacBio reads with Flye (v.2.4). Contigs were then polished using the Illumina and ONT reads. For the mitogenome, a reference-based approach was followed. The mitogenome sequence of *Hanseniaspora uvarum* (DQ058142) was downloaded from NCBI and PacBio reads were mapped to the reference using minimap2 (set parameters: -a, -x map-pb). Mapping reads were filtered from the bam file using samtools (-F 4) and extracted into a fastq file using bam2fastq (v1.1.0). The reads were assembled using Flye (v.2.4; default parameters, except: estimated genome size of 20 kb (40)), which resulted in the assembly of the 17 kb linear mitogenome. PlasmidSpades (41), run on the Illumina data, did not detect plasmids. The mean telomere length (pattern “CCTGA”) was calculated using the Illumina reads and computel (v.1.2; (42)) and was estimated to be 2275 bp. The number of telomere patterns at both ends of each contig was counted manually (Supplementary Table S1). PloidyNGS (v.3.1.2; (43)) as well as nQuire (44) estimated the genome to be diploid. The extensive polishing and manual curation resulted in a total of 7 chromosomes and 1 mitogenome. The total genome size was 8,767,711 bp.

*H. meyeri* (APC 12.1) genes were identified as previously described by using the Yeast Genome Annotation Pipeline (YGAP) (38). KOALA was used to assign KEGG Orthologs (KOs; K numbers) to the predicted proteins (45). The KEGG Mapper Reconstruct tool was used to assign the KOs to pathway modules (46). The genome annotation and all sequencing data are available at the Harvard Dataverse (https://doi.org/10.7910/DVN/PXBEDO) and NCBI under the bioproject PRJNA907227.

### Cloning and transformation

Pantothenate transporter genes (*PANT1-6*) were synthesized *in silico* (Twist Bioscience, South San Francisco, USA; codon optimised for *S. cerevisiae* and lacking the restriction sites Bsa I, BsmB I, and Not I) and introduced into expression vectors using golden gate cloning ((47); MoClo Yeast Toolkit, Addgene, Watertown (MA), USA). Constitutive expression of each transporter was achieved with the assembly of pTDH3 promoter and tTDH1 terminator modules into a GFP drop-out vector. The reaction was performed in temperature cycles of 37°C for 2 min and 16°C for 5 min using the restriction enzyme Bsa I-HF® V2 and the Hi-T4™ DNA ligase (from New England Biolabs; Bioconcept, Allschwil, Switzerland), followed by a final 10 min ligation at 16°C. Intermediate and final constructs were transformed into *E*.*coli* DH5α and purified with the Qiaprep Spin Miniprep Kit (Qiagen, Hombrechtikon, Switzerland). The size of the final products was confirmed by restriction digestion with Not I and gel electrophoresis.

The *FEN2* promoter of *Δpan6 S. cerevisiae* mutants was exchanged by chromosomal integration of a pGAL.L-containing PCR product (48) to construct the reporter strain. Briefly, primers were designed with homology to the *FEN2* promoter region for amplification of promoter GAL.L using the plasmid pYM-N27 from the PCR toolbox (Euroscarf/Scientific Research & Development GmbH, Oberursel, Germany)(48). Cells were selected in YPD with nourseothricin (100 μg/mL; Jena Bioscience, Jena, Germany) after incubation at 30°C for 3 days. All primers used in this study are listed in Supplementary Table S3.

Golden gate constructs (47) containing *PANT1-6* genes, were linearized with Not I for integration into the *URA3* locus of the *S. cerevisiae* genome. After lithium acetate transformation, cells were grown in yeast nitrogen base without histidine and with galactose as carbon source

### Fluorescence microscopy

Cells were grown overnight at 30°C in YPD medium, diluted to OD_600_ 0.1 and harvested when the culture reached 0.5. Cells were spread on slides coated with an SC agar patch. Stacked images were recorded at a spinning disc confocal inverted microscope (Zeiss Axiovert 200m) using the 100x oil objective and an Evolve EMCCD Camera (Photometrics).

## Acknowledgments

We thank Maja Hilber-Bodmer and Laurin Müller for support in the laboratory and Antonio Di Pietro for providing the *Fusarium oxysporum* f. sp. *lycopercisi* strain. Many helpful comments and suggestions from Kenneth H. Wolfe are also greatly appreciated. Jürg Frey, Daniel Frei, and Inés Sumann are acknowledged for generating ONT and Illumina data.

## Notes

https://dataverse.harvard.edu/dataverse/Hans_genome

### Competing Interest Statement

The authors have declared no competing interest.

