## Supplementary material for "Pantothenate auxotrophy in a naturally occurring biocontrol yeast"

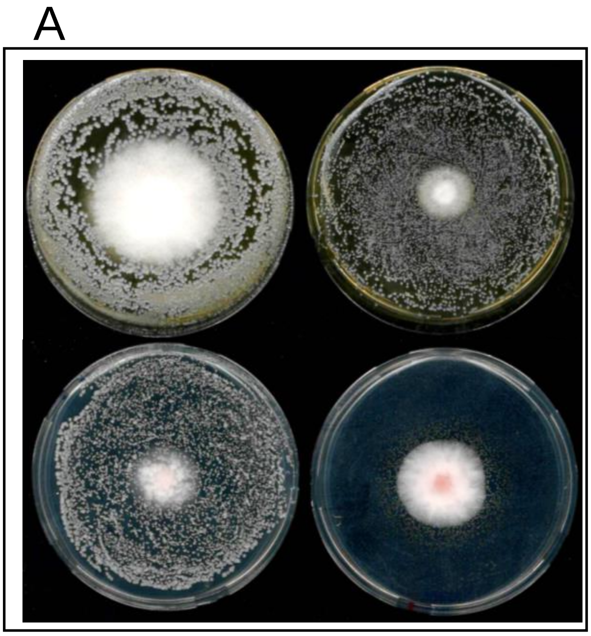


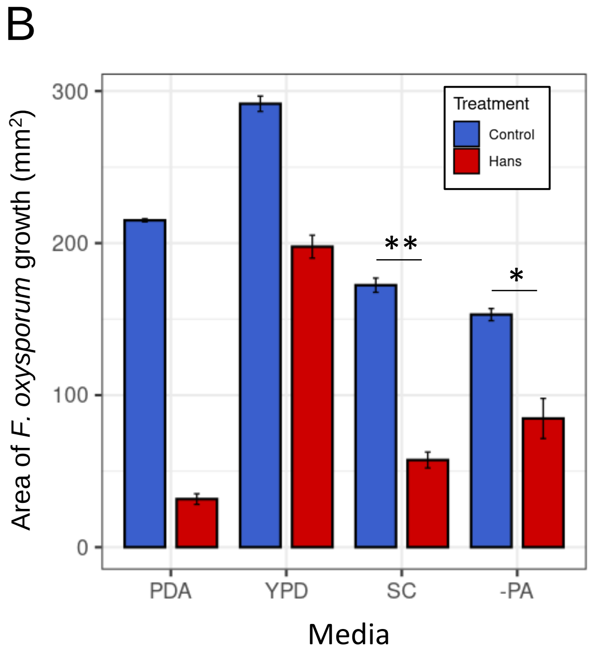


**Supplementary Figure 1**

**A**. Competition between *Hanseniaspora meyeri* and *F. oxysporum* f. sp. *lycopersici* in YPD, PDA (top row), SC and SC without pantothenate (-PA, bottom row). The yeast grows on the surface of the plate in the first three media, but not in the absence of pantothenate. The *F. oxysporum* colony is larger in YPD, but growth is reduced in all media. **B**. Control of *F. oxyporum* was the strongest in PDA, the area was reduced to 15% of the growth area without *H. meyeri*. In SC media, with and without pantothenate, *Fusarium* growth is reduced to 33% and 55%. p.adj FDR method -PA*: 0.0270000, SC: **0.0001776

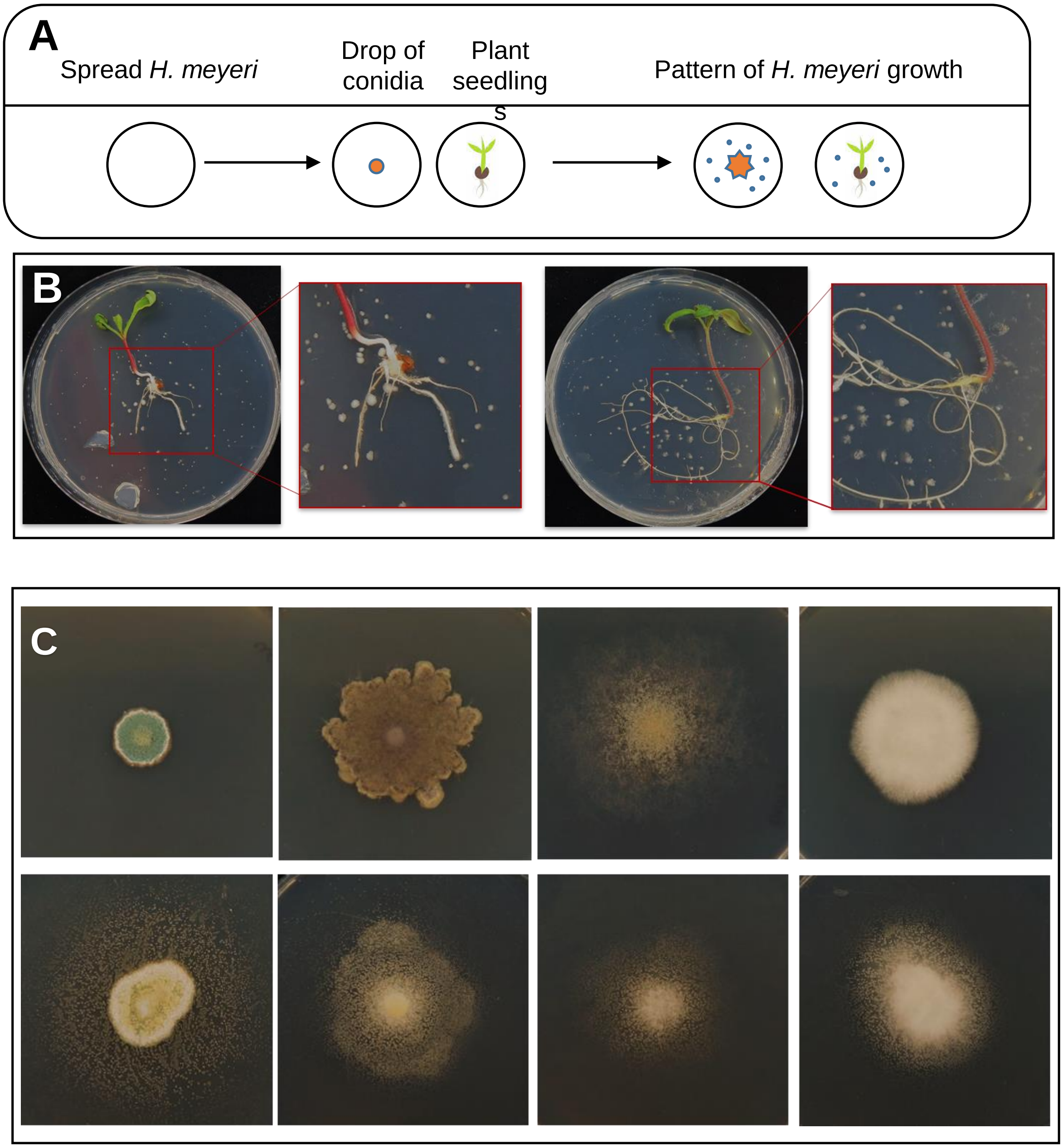


**Supplementary Figure 2**

**A.** Diagram of co-culture assays between *H. meyeri* APC 12.1 and filamentous fungi or plant seedlings in MS media **B.** Coculture with radish and tomato plants and **C.** competiton assays with four plant pathogenic fungi in Murashige-Skoog media shows the yeast grows only in the vicinity of other organisms. **Top row** : Filamentous fungi colony growing alone, **Bottom row** : *H. meyeri* (APC 12.1) and filamentous fungi competition assay. From right to left: *Penicillium polonicum, Mucor moelleri, Botrytis caroliniana, Fusarium oxysporum.*

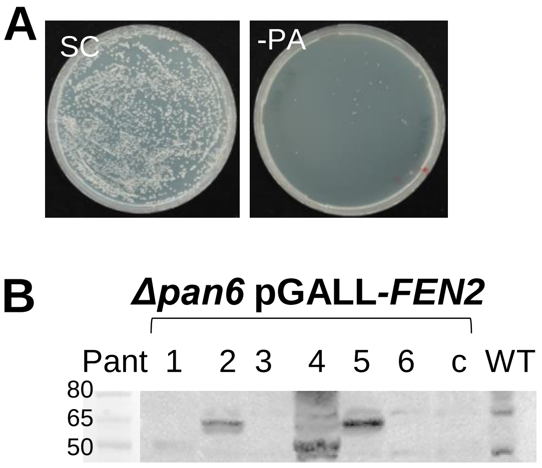


**C**

| Protein | Expected size (kDa) |
|---|---|
| Pant1 | 63.57 |
| Pant2 | 67.9 |
| Pant3 | 72.28 |
| Pant4 | 63.03 |
| Pant5 | 74.37 |
| Pant6 | 62.67 |

**Supplementary Figure 3**

**A.** Phenotype of *S. cerevisiae* BY4741 Δ*pan6* (Y02304) in SC and SC without pantothenate. **B.** Expression of 3xFLAG-6xHis tagged Pant2, Pant4 and Pant5 was confirmed by anti-6xHis western blot. Pant1, Pant3 and Pant6 do not seem to be expressed. **C.** Expected sizes of protein products for each tagged pantothenate transporter.
